# MECOM is a master repressor of myeloid differentiation through dose control of *CEBPA* in acute myeloid leukemia

**DOI:** 10.1101/2025.01.02.631046

**Authors:** Dorien Pastoors, Marije Havermans, Roger Mulet-Lazaro, Leonie Smeenk, Sophie Ottema, Claudia Erpelinck-Verschueren, Stanley van Herk, Maikel Anthonissen, Tim Grob, Bas J. Wouters, Berna Beverloo, Torsten Haferlach, Claudia Haferlach, Johannes Zuber, Eric Bindels, Ruud Delwel

## Abstract

Enhancer translocations, due to 3q26 rearrangements, drive out-of-context *MECOM* expression in an aggressive subtype of acute myeloid leukemia (AML). Direct depletion of MECOM using an endogenous auxin-inducible degron immediately upregulates expression of myeloid differentiation factor *CEBPA*. MECOM depletion is also accompanied by a severe loss of stem cells and gain of differentiation. MECOM exerts its inhibitory effect by binding to the +42kb *CEBPA* enhancer, a gene essential for neutrophil development. This is partially dependent on the interaction between *MECOM* and its co-repressor CTBP2. We demonstrate that *CEBPA* overexpression can bypass the MECOM-mediated block of differentiation. In addition, AML patients with *MECOM* overexpression through enhancer hijacking show significantly reduced *CEBPA*. Our study directly connects two main players of myeloid transformation *MECOM* and *CEBPA*, and it provides insight into the mechanism by which MECOM maintains the stem cell state in a unique subtype of AML by inactivating *CEBPA*.

## Introduction

Activation of the *MECOM* (*MDS1* And *EVI1* Complex Locus) locus at chromosome 3q26 in healthy bone marrow is mostly restricted to hematopoietic stem and progenitor cells (HSPCs) ^1^. *MECOM* encodes for two main isoforms: a short isoform, *EVI1*, a zinc-finger transcription factor that is required for hematological development and maintenance of hematopoietic stem cells ^2,3^. The long isoform (*MDS1-EVI1*) contains two additional exons with putative methyltransferase activity. *MECOM* was first identified as a mouse retroviral insertion site, where insertion of proviral DNA near *Evi1* causes its constitutive transcriptional activation, leading to leukemia ^4,5^. A similar out of context over-activation of *MECOM* occurs because of enhancer translocations or inversions involving chromosome 3q26 in AML. For instance, in AMLs with inv(3)(q21q26)/t(3;3)(q21;q26), a distal enhancer of *GATA2* translocates to *MECOM* to drive *EVI1* (not *MDS1-EVI1*) overexpression ^6–9^. As the official gene symbol is *MECOM*, we will use *MECOM* to denote *EVI1* expression in 3q26-rearranged AML. These enhancer hijacking events activate *MECOM* beyond the restricted cell state where it is active in healthy bone marrow. Its expression is essential for the survival and immature phenotype of those AML cells^8,10,11^. It is important to note that AMLs with 3q26/*MECOM* rearrangements exhibit a very poor therapy response and low survival rates ^12–15^.

MECOM has been reported to be a context-dependent gene repressor and activator, and many putative up- and downregulated target genes have been identified. MECOM has been reported as a positive regulator of *MiR1-2*^16^, *Spi1*^17^, *Fbp1*^18^*, Erg*^19^, *NRIP1*^20^ and *PTGS2*^21^. Examples of genes reported to be negatively regulated by MECOM are *miR-449A*^22^, *Serpin B2*^23^, *miRNA-124*^24^, *RUNX1*^25^, *PTEN*^26^ and *CEBPA*^27^. MECOM has also been described as a repressor for TGFB signaling^28,29^ and NFκB^30^, as well as an activator of NOTCH^31^. Thus, although many different MECOM target genes have been proposed, none of them have been found recurrently. Therefore, we set out to uncover critical gene(s) involved in MECOM-driven leukemia development and block of myeloid differentiation.

Here, we studied the effects of rapid MECOM degradation on gene regulation, using an auxin-inducible MECOM degron system. Upon auxin treatment, MECOM is degraded within 1.5 hours, enabling rapid profiling of newly made RNA (SLAM-Seq), and of chromatin (ATAC-seq, ChIP-seq and 4C). Using this system, we show that MECOM directly represses *CEBPA* and this is crucial for the differentiation block of inv(3) leukemia cells. This repression happens via binding to the +42kb *CEBPA* enhancer which is indispensable for neutrophil development^32,33^. Our study provides insight into the mechanism of stem cell maintenance by *MECOM* in a unique AML subtype.

## Results

### Degradation of MECOM drives myeloid differentiation of inv(3)/t(3;3) AML cells

To rapidly deplete MECOM protein, we introduced an auxin-inducible degron (AID) (V5-AID-T2A-eGFP) 3’ of *MECOM* in the inv(3) cell line MUTZ3 (Fig 1A, Fig S1A) ^34^. The cells were stably transduced with an OsTIR1 construct, required for tethering the AID tag to the nuclear ubiquitination complex in presence of auxin (Fig S1A). Three separate clones were derived for subsequent experiments. Upon auxin exposure, MECOM protein is degraded within 1.5 hrs (Fig 1B, Fig S1B, Fig S1C). When cells were cultured for seven days with auxin-supplemented media (refreshed daily, Fig S1B), they lost CD34 and gained CD15 cell surface markers, indicative of myeloid differentiation (Fig 1C, Fig 1D). Interestingly, the cells in the CD34-CD15+ fraction were eGFP negative, even though GFP was not degraded by auxin because it is separated from MECOM-AID by a T2A self-cleavage site (Fig 1E). This suggests that, when MECOM is eliminated, cells differentiate and lose the ability to transcribe *MECOM* and thus *GFP* driven by the hijacked *GATA2* enhancer.

**Figure 1:**
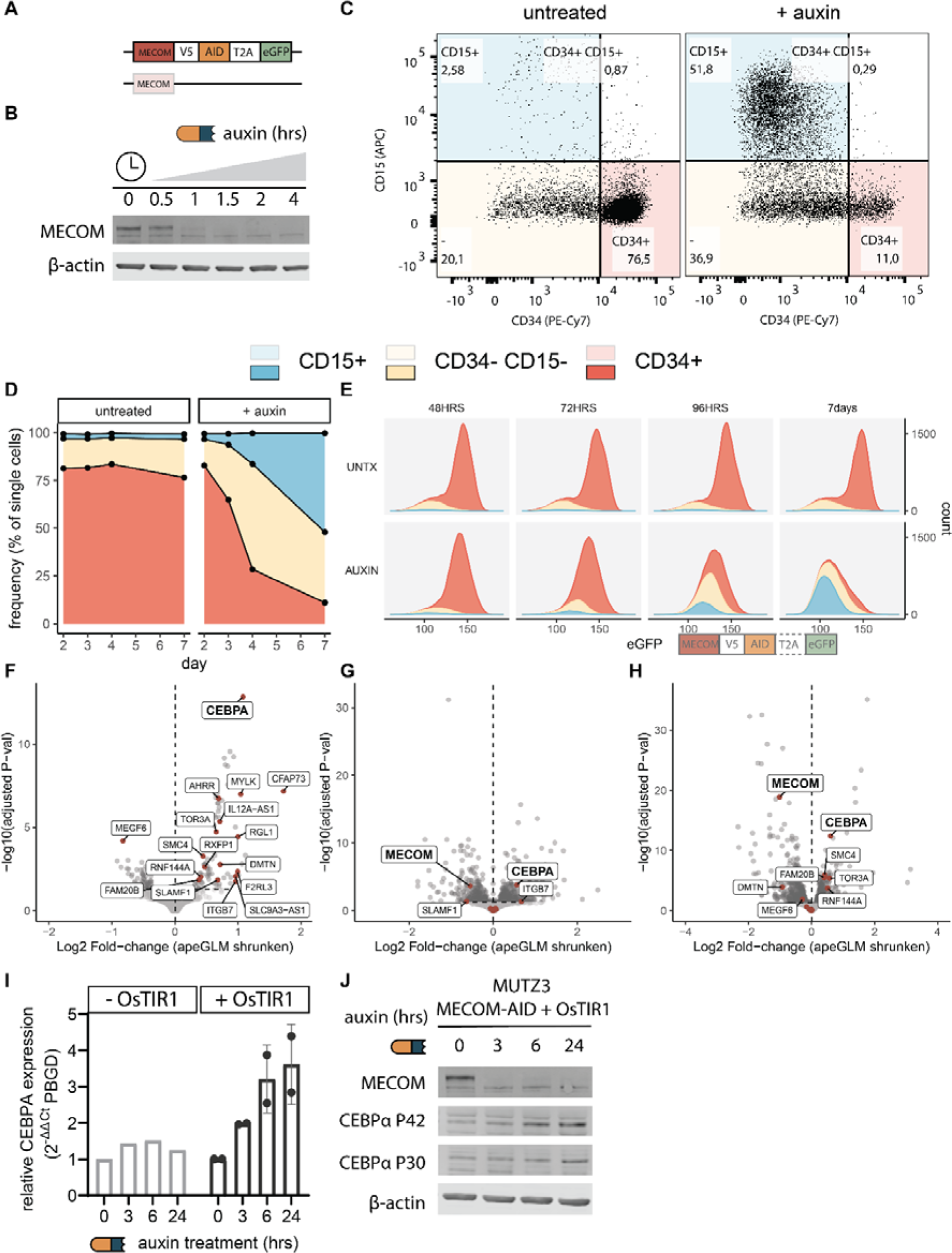
*CEBPA* is a target of MECOM. Fig 1A. Schematic depiction of the AID tag, followed by V5-T2A-eGFP introduced 3’ of MECOM in MUTZ3 cells. Fig 1B. Western blot analysis of a MUTZ3 AID-TIR1 clone for MECOM (upper panel) and B-actin (lower panel). Cells were treated with 500 µM of auxin for indicated time spans. Fig 1C. Flow-cytometric analysis of CD34-PeCy7 and CD15-APC staining in a MUTZ3 AID-TIR1 clone treated with auxin (100 µM, refreshed every 24 hours) after 7 days of auxin treatment. Fig 1D. Flow-cytometric analysis of a single MUTZ3 AID-TIR1 clone treated with auxin (100 µM, refreshed every 24 hours) for 7 days. Frequencies reported as percentage of single-cells based on the gates shown in panel C. Fig 1E. Stacked density histogram proportional to counts per group of eGFP in MUTZ3-AID T2A eGFP cells treated with auxin. Gates and legend are shared with 1C and 1D. Fig 1F. Volcano plot of DEseq2 analysis on T>C-converted reads (nascent RNA) (auxin versus no auxin), on three MUTZ3 AID OsTIR1 clones from panel B. Genes are labelled if the adjusted p-value is below 0.01 and they are specifically upregulated in MUTZ3-MECOM-AID cells treated with auxin, and not in auxin-treated WT cells. Fig 1G. Volcano plot of DEseq2 analysis on MUTZ3 treated with two shRNAs targeting MECOM versus a scrambled control after 24 hrs of puro selection (started 48 hrs post-transduction) (n=3 for each shRNA, 2 shRNAs directed against MECOM versus 1 shRNA_control). These two shRNAs downregulate MECOM (as shown in plot). Genes from panel F are labelled in this plot if the adjusted p-value is below 0.05. Fig 1H. Volcano plot of DEseq2 analysis on MUTZ3 expressing doxycycline-inducible Cas9, treated with 3 guides inactivating a GATA2 enhancer in control of MECOM (doxycycline versus no doxycycline). Previous work^10^ has shown these three guides downregulate MECOM (as shown in plot). Genes from panel F are labelled in this plot if the adjusted p-value is below 0.05. Fig 1I. Bar plot showing qPCR of *CEBPA* (normalized to PBGD, gene symbol *HMBS*) of MUTZ3 AID-TIR1 clone treated with 80 µm auxin at indicated timepoints. Fig 1J. Western blot analysis of MUTZ3 AID-TIR1 clone treated with 80 µm auxin at indicated timepoints.

### *CEBPA* is rapidly activated upon degradation of MECOM in inv(3)/t(3;3) AML

To identify direct target genes of MECOM we performed metabolic labelling of newly produced RNA (SLAM-Seq) upon degradation of MECOM. After 1.5 hours of auxin treatment, the MECOM-AID expressing MUTZ3 cells were exposed to uracil analogue S4U for three hours leading to T>C conversion in sequencing reads derived from RNA produced during this time frame (Fig S1B) ^35^. Differential gene expression analysis on T>C converted RNA revealed seventeen differentially expressed (DE) genes, of which sixteen were upregulated upon MECOM elimination (Fig 1E, Fig S1C, Fig S1D, Fig. S2A, Fig S3A). Of those seventeen DE genes, *CEBPA* is the only one that is consistently upregulated when *MECOM* is knocked-down using two distinct short hairpin RNAs (shRNAs) in MUTZ3 cells (Fig 1G). Inactivating mutations introduced by CRISPR/Cas9 in the hijacked *GATA2* enhancer driving MECOM expression^10^ were also accompanied by *CEBPA* upregulation (Fig 1H). *CEBPA* levels increased further after 3, 6 and 24 hrs upon auxin exposure in OsTIR1-containing MUTZ3 cells (Fig 1I). An increase of CEBPA protein was evident at 6 and 24 hours after MECOM elimination (Fig 1J). Taken together, these data indicate that *CEBPA* expression is rapidly and persistently upregulated upon MECOM downregulation.

### MECOM degradation changes chromatin state in the *CEBPA* locus

ATAC-seq in MUTZ3 MECOM-V5-AID OsTIR1 clones revealed a small set of cis-regulatory elements (CRE) that gained accessibility 24 hours after MECOM degradation. Most of those sites were unchanged 4 hours after MECOM depletion (Fig 2A and Fig 2B). Strikingly, the most significantly changed CRE was the previously identified +42kb enhancer, which is indispensable for *CEBPA* transcription in myeloid progenitors and neutrophil development ^32,36,37^ (Fig 2B, Fig 2C, Fig S4A). MECOM is indeed bound to the +42kb enhancer of *CEBPA* in MUTZ3 and primary AML cells with an inv(3) rearrangement as shown by ChIP-sequencing (Fig 2C). We have previously characterized components of the MECOM protein complex, which consists of corepressors CTBP1/2 and other repressive proteins, including H3K9- and H3K27 methylating enzymes ^11^. Accordingly, MECOM depletion was accompanied by an almost complete loss of binding of corepressor CTBP2 to the +42kb *CEBPA* enhancer (Fig 2C). In fact, loss of CTBP2 binding was observed at all binding sites in the genome upon MECOM depletion (Fig. S5C). In addition, MECOM loss was accompanied by a minor increase of RUNX1 and P300 binding at the +42 kb enhancer (not significant) (Fig 2C and S2B). H3K27Ac levels significantly increased at the +42kb enhancer (Fig 2C and Fig S4A), and also at the +55kb and +34kb CREs (Fig 2C and Fig S4A). In line with increased *CEBPA* transcription, H3K27ac levels were also increased at the *CEBPA* gene body upon MECOM depletion (Fig 2C and Fig S4A). Loss of MECOM did not cause a change in promoter-enhancer interaction in the locus of *CEBPA* (Fig S4A). Repressive marks H3K27Me3 and H3K9Me3 were unchanged at this region (Fig S4A) as well as globally (Fig S4A, Fig. S5A, Fig. S5B). Together our data are in line with a repressive function of MECOM on *CEBPA* transcription by binding to the +42KB enhancer.

**Figure 2:**
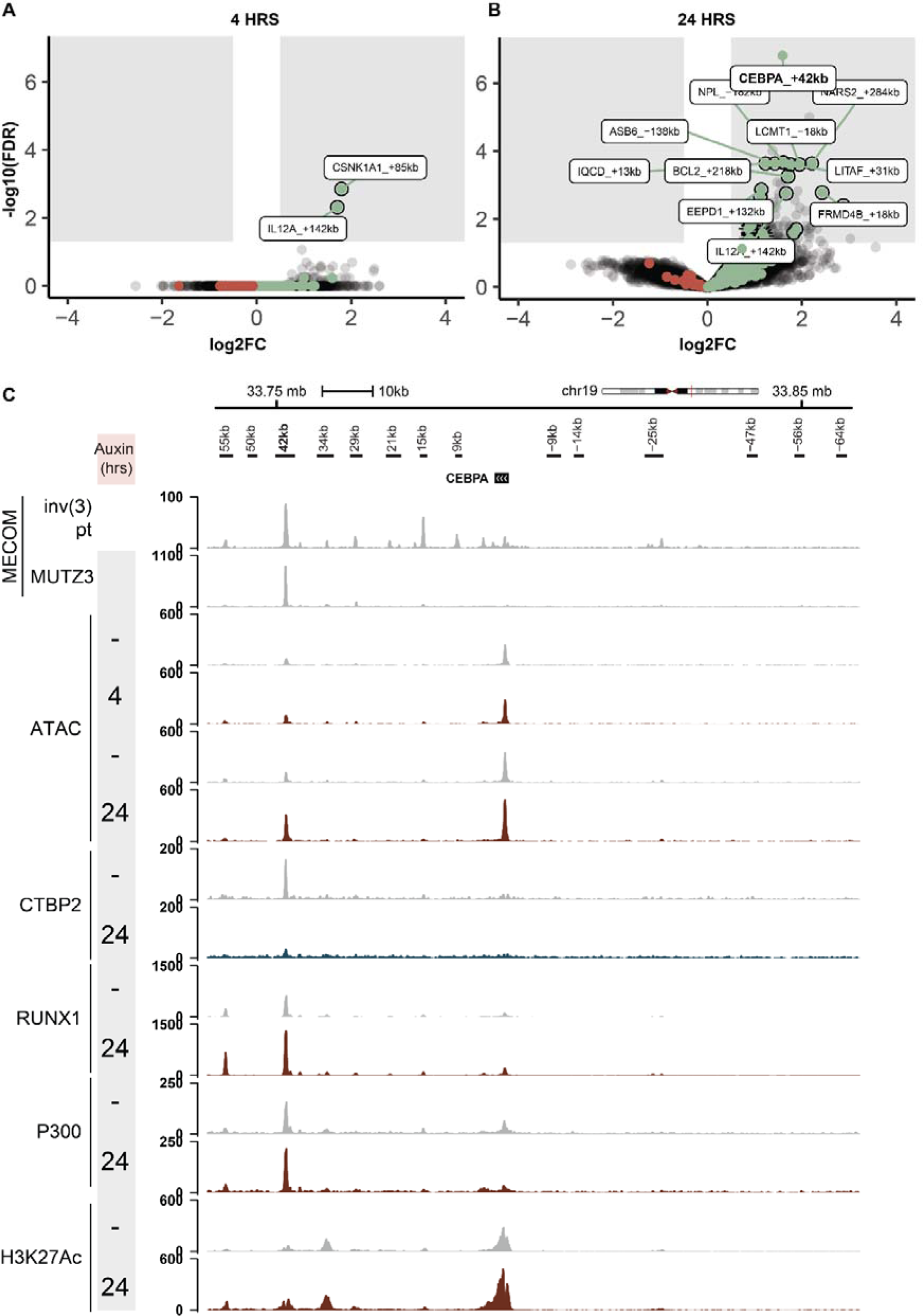
MECOM regulates CEBPA through its +42 kb enhancer. Fig 2A. Volcano plot of differential chromatin accessibility (ATAC-seq) of untreated (n = 3) versus auxin-treated for 4 hrs (n = 2; 500 µM) calculated with DiffBind. Fig 2B. Volcano plot of differential chromatin accessibility (ATAC-seq) of untreated (n = 3) versus auxin-treated for 24 hrs (n = 2; 500 µM) calculated with DiffBind. All labels containing “CEBPA” at FDR < 0.05 are labelled in bold. Fig 2C. ATAC-seq and ChIP-seq for indicated factors in unmodified cells (MECOM tracks; inv(3) patient and MUTZ3 bulk) and a MUTZ3 MECOM-V5-AID OsTIR1 clone (ATAC, RUNX1, CTBP2, P300, H3K27ac), untreated and auxin-treated for indicated timepoints (500 µM). Coordinates are hg19 and tracks are RPKM normalized.

### Depletion of MECOM activates the CEBPA regulatory program in 3q26-rearranged AML cells

RNA-seq in MUTZ3 cells 24, 48, 72, 96 and 120 hrs after *MECOM* knock-down revealed significant *CEBPA* upregulation at all timepoints (Fig. 3A, Fig. S6A). GSEA showed that genes known to be activated by CEBPA are upregulated in a time-dependent manner following *MECOM* knock-down (Fig. S6B). Besides the +42kb enhancer of *CEBPA* (Figure 2B), multiple other CREs became more accessible upon MECOM depletion. These may include direct as well as indirect MECOM targets. To examine which factors might drive these secondary effects, we carried out transcription factor footprinting which predicted an increase in binding of CEBP factors to their motifs (Fig. 3A). While this change in CEBP motif protection was already significant 4 hrs post MECOM depletion, the effect increased at 24 hrs (Fig 3A, Fig. S6C, Fig S7A). Although less prominent, we also observed gain of protection of NFIL sites and loss of FOSL1 motif binding. We also performed differential motif enrichment analysis on our ChIP-seq data. In all upregulated H3K27ac, P300 and RUNX1 peaks, CEBP motifs are highly enriched (Fig. 3D, Fig. 3E, Fig S7B,Fig S7C). In addition, we found enrichment of primarily ETS-like motifs in the downregulated peaks where significant enrichment is found (Fig. 3D, Fig. 3F, Fig S7B,Fig S7C). Together those analyses show that MECOM has a major repressive effect on the CEBPA regulatory program, and that induction of *CEBPA* expression has profound impact on the chromatin- and expression landscape in t(3;3)/inv(3) AML.

**Figure 3.**
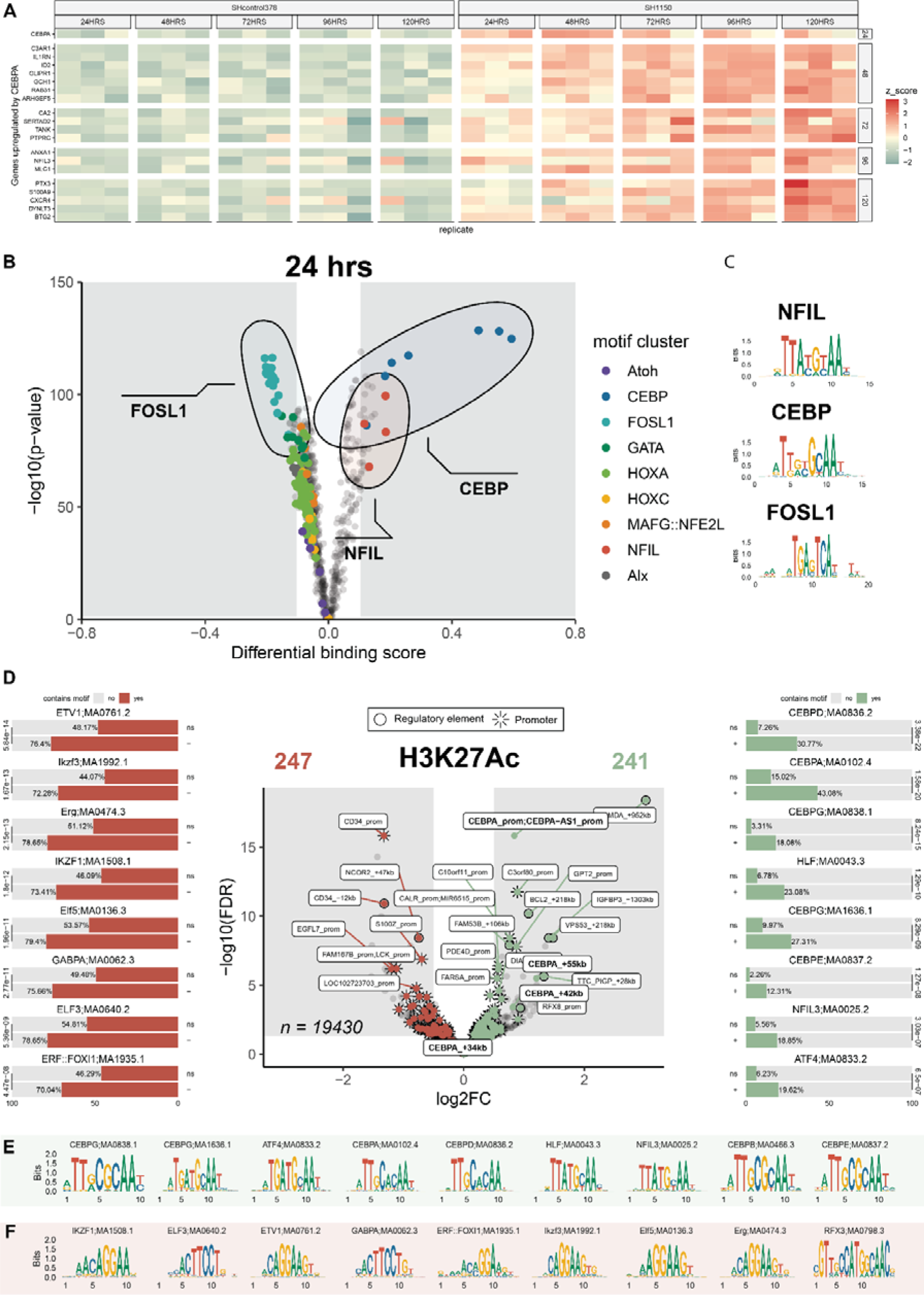
*CEBPA* activation has a major effect on chromatin after MECOM depletion. Fig. 3A. Heatmap of z-scores of RNA-seq rlog-transformed counts of MUTZ3 cells treated with control shRNA or MECOM shRNA. Only genes previously known to be upregulated by *CEBPA* are shown^47^, grouped according to the first time point at which they are upregulated following MECOM KD. From rlog transformed counts, the mean of the control samples is subtracted prior to z-score calculation. This heatmap shows 1 of the two shRNAs; the other is in Fig. S6A. Fig. 3B. Volcano plot ATAC-seq footprinting of on MUTZ3-V5-AID-OsTIR1 cells after 24 hrs of auxin depletion (n = 2) compared to untreated control (n = 4). Each dot is a single motif, and motifs are colored by their cluster. Analysis performed with TOBIAS with aggregated reads in each group. Fig. 3C. Composite motifs of the indicated motif clusters from panel A. Fig. 3D. Volcano plot of differential H3k27Ac signal (ChIP-seq) in MUTZ3-V5-AID-OsTIR1 cells after 24 hrs of auxin depletion versus untreated control (n = 2). A list of peaks was compiled based on differential binding across experiments as detailed in the Methods. AME motif enrichment is shown on the corresponding side of the plot. Bars represent the fraction of peaks that contain a high-score motif per set; significance is indicated with AME e-value across the bars. Top-8 motifs are shown per ChIP at an e-value cut-off of 0.05. All labels containing “CEBPA” at FDR < 0.05 are labelled in bold. Fig. 3E. Motif logos of all unique motifs found in motif enrichment analysis with AME in upregulated peaks of any factor. Fig. 3F. Motif logos of all unique motifs found in motif enrichment analysis with AME in downregulated peaks of any factor.

### The +42kb *CEBPA* enhancer is essential to induce differentiation when MECOM is eliminated

As CEBPA is a master regulator of myeloid differentiation, our findings suggest that the +42kb enhancer is important for the onset of differentiation after MECOM is depleted. To test this hypothesis, we generated +42kb *CEBPA* enhancer-deleted MUTZ3 clones (Fig. 4A). While we screened >100 clones, we were only able to derive 42kb^Δ/+^ clones. Upon depletion of MECOM levels with a sgRNA directed to MECOM, myeloid differentiation (CD15+ CD34-) was no longer induced in the 42kb^Δ/+^ clones while it was in the 42kb^+/+^ cells (Fig. 4B,Fig. S8A). Western blot analysis showed that CEBPA was only marginally upregulated in the 42kb^Δ/+^ clone (Fig. 4C). This suggests a dose-dependent effect of CEBPA on differentiation, where a certain threshold-amount of CEBPA is required to efficiently induce differentiation. This also explains the differentiation block despite low-level *CEBPA* expression in these cells.

**Figure 4:**
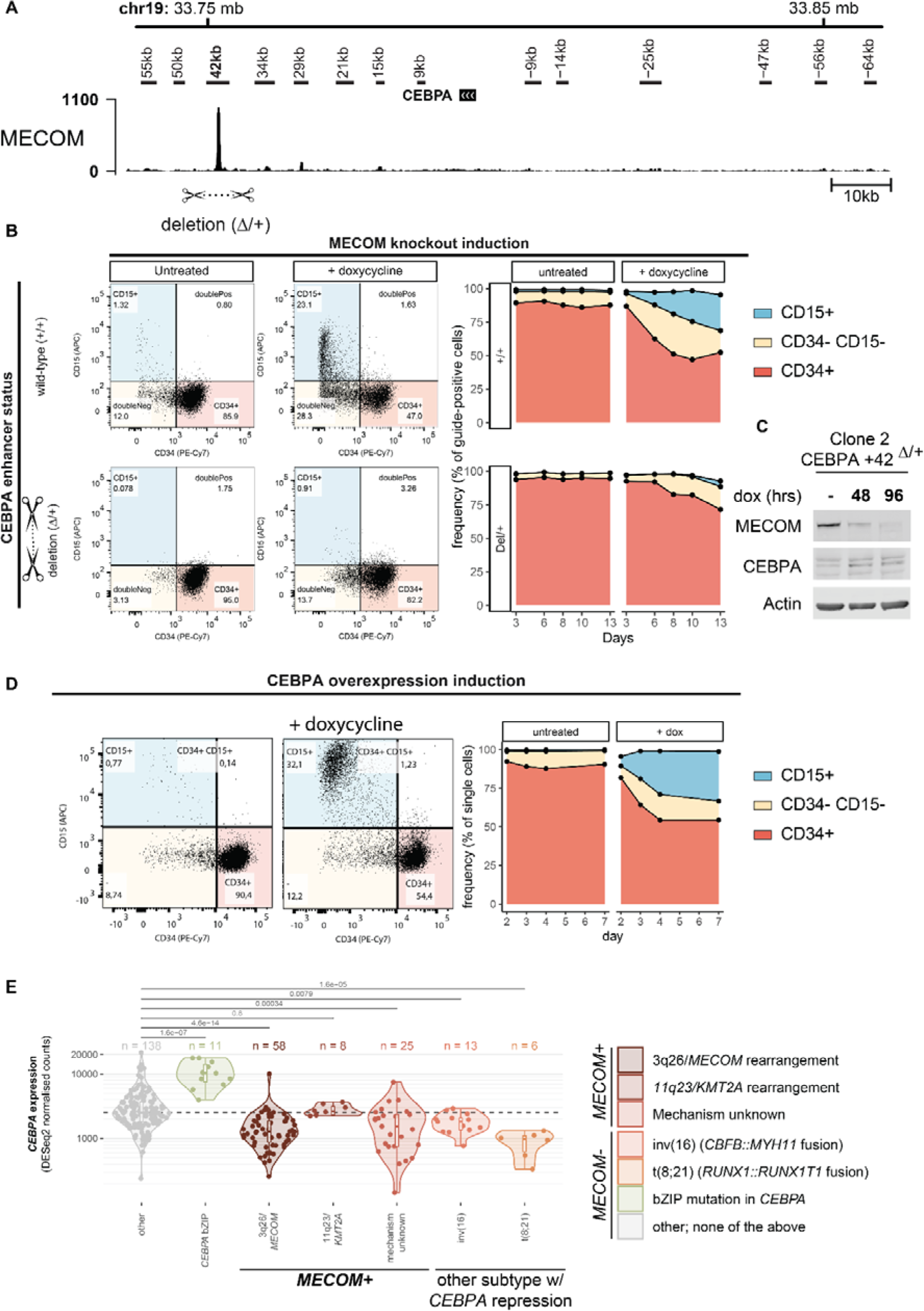
C*E*BPA is repressed by MECOM in MUTZ3 and in primary AML samples. Fig. 4A. ChIP-seq for MECOM in the *CEBPA* locus with the deletion of the +42kb enhancer used in panels B and C is indicated. The panel with MECOM is the same as in Fig. 2C. Coordinates are hg19 and data is RPKM normalized. Fig. 4B. Differentiation of MUTZ3 MECOM-eGFP iCas9 upon MECOM knockout in wild-type (top) and a heterozygous clone (bottom; deletion of +42kb enhancer indicated below panels A and B). Day 10 shown (left panels) and days 3-13 (right panels). Fig. 4C. Western blot of MECOM and CEBPA in a MUTZ3 clone with heterozygous deletion of the +42kb *CEBPA* enhancer. Fig. 4D. Flow-cytometric analysis of MUTZ3 cells transduced with a lentiviral dox-inducible *CEBPA* construct, treated with doxycycline (1 ug/mL) for 7 days and stained with CD15-APC and CD34-PeCy7 (day 7 shown). Flow-cytometric analysis from panel A, shown at days 2-7. Frequencies reported as percentage of single cells based on the gates in shown in panel B (days indicated). Fig. 4E. Normalized *CEBPA* expression in AML subtypes with MECOM or *CEBPA* involvement, compared to controls (DESeq2-normalized counts). A case with a complete CEBPA deletion, as well as CpG Island Methylator Phenotype cases with heavily methylated *CEBPA* promoter were excluded from the analysis due to their complete lack of expression of *CEBPA*. Adjusted P-values from DESeq2 are reported for indicated comparisons and group sizes are indicated above each boxplot. A dashed line indicates the median of the control group.

### *CEBPA* overexpression induces differentiation in the presence of MECOM

If increased CEBPA expression upon MECOM degradation is the main driver of differentiation, overexpression of *CEBPA* should bypass the differentiation block enforced by MECOM. Doxycycline-induced *CEBPA* overexpression in MUTZ3 cells indeed strongly increased the fraction of CD15+ cells, while reducing the fraction of CD34+ cells (Fig. 4D). Taken together, our results suggest *CEBPA* is repressed by MECOM in AML with 3q26/*MECOM* rearrangements.

### CEBPA mRNA levels are significantly reduced in 3q26-rearranged AMLs

We carried out RNA-seq in 58 bone marrow aspirate or peripheral blood samples from newly diagnosed AML patients with 3q26/*MECOM* rearrangements. This cohort contained 12 cases with inv(3)/t(3;3) and 46 cases with alternative 3q26 rearrangements. Differential gene expression revealed that AML samples with 3q26 rearrangements express significantly reduced *CEBPA* mRNA levels compared to non-3q26 rearranged samples (Fig. 4E)^6,38–40^. This repressive effect was similar in magnitude in samples obtained from AML patients with with inv(16) (*CBFB::MYH11* fusion) or t(8;21) (*RUNX1::RUNX1T1*), which are known to have low *CEBPA* expression^41,42^. In case of AMLs with a translocation t(8;21) it has been proposed that the RUNX1-RUNX1T1 fusion protein represses CEBPA transcription via the +42kb enhancer as well^42,44,45^. We conclude that repression of *CEBPA* transcription is an essential event in leukemic transformation by MECOM in 3q26-rearranged AML.

### *CEBPA* is partially upregulated when MECOM-CTBP interaction is blocked

We have previously developed a 4xPLDLS overexpression peptide construct which is able to outcompete MECOM-CTBP2 interaction^11^. This construct fully inhibits in vivo leukemia development, indicating that MECOM-CTBP2 interaction is essential for leukemic transformation of *MECOM* rearranged AML. To determine if repression of *CEBPA* transcription by MECOM is dependent on CTBP2, we derived 4 inducible 4xPLDLS clones from MUTZ3 cells (Fig S9A). Upon induction of 4xPLDLS with doxycycline, we observed a complete loss of CTBP2 repressor binding to the +42kb enhancer (Fig 5A), accompanied with a predominant upregulation of *CEBPA* among other genes (Fig 5B, Fig 5AC). We observed similar *CEBPA* upregulation in MUTZ3 cells transduced with a 4 x PLDLS retroviral construct compared to a 4x PLASS control^11^ (Fig S9B, Fig 5CD, Fig 5DE). Flowcytometric analysis revealed a consistent increase of CD34-/CD15+ differentiated cells when 4 x PLDLS was expressed over a prolonged time, in contrast to 4 x PLASS expressing M<UTZ3 cells (F). Positive enrichment of upregulated CEBPA target genes (Fig S9C) emphasizes a role for CTBP in the repression of *CEBPA* by MECOM.

**Figure 5:**
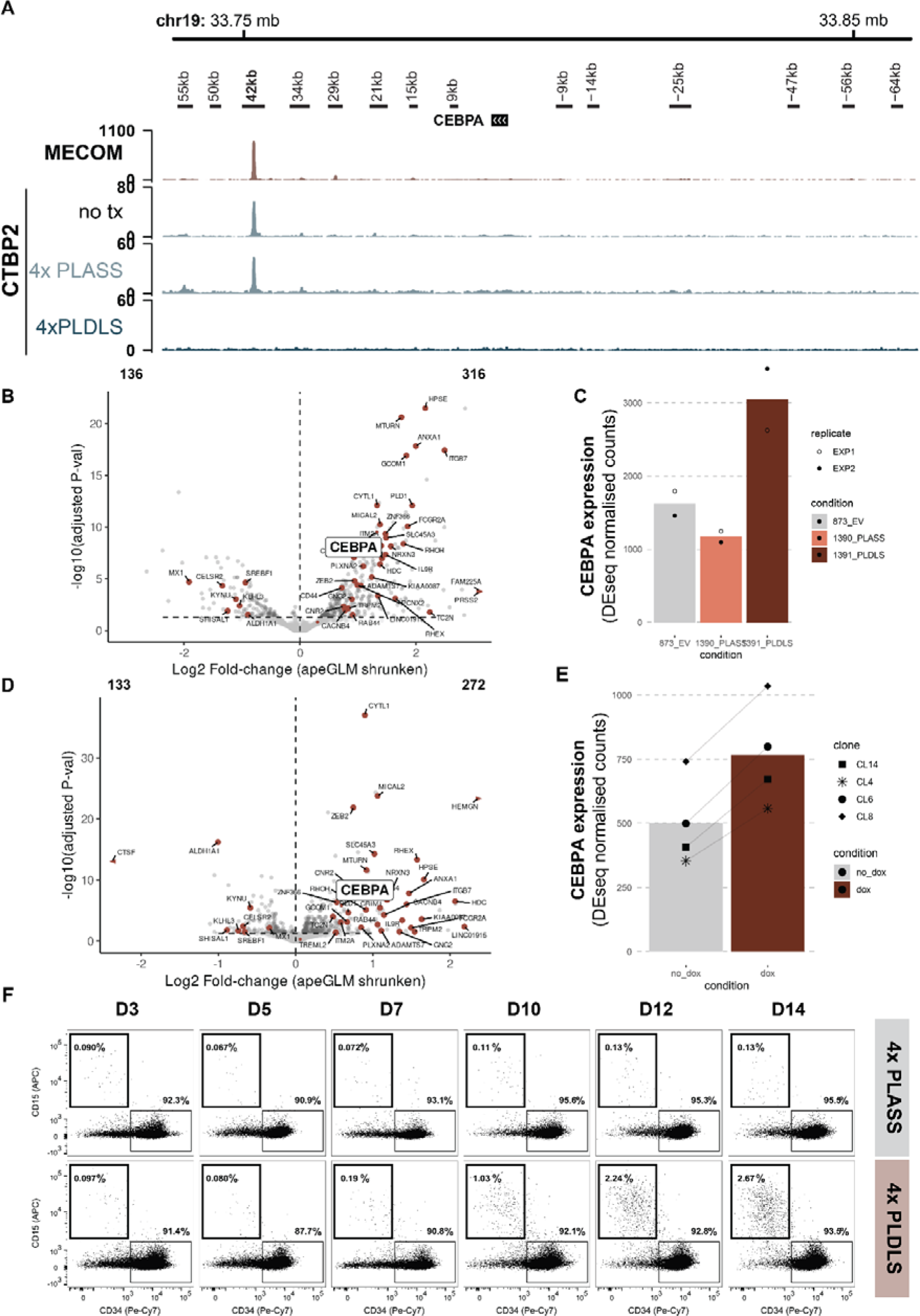
C*E*BPA is partially de-repressed upon induction of MECOM-CTBP2 interaction inhibitor peptide. Fig 5A. Volcano plot of DEseq analysis of 4 doxycycline-treated pCW-4xPLDLS clones versus untreated (72hrs; design: ∼ clone + condition). The horizontal line indicates a significance level of padj < 0.05. Arrows indicate genes outside axis limits. Top-40 consistently changed genes between A and C are labelled (total overlapping: down = 22, up = 70). Fig 5B. Barplot of DESEQ normalized counts for *CEBPA* (Experiment as in A.) Fig 5C. Volcano plot of DEseq analysis of 2 replicate experiments in MUTZ3 of 4xPLDLS transduced versus 4x PLASS (72hrs of selection; design: ∼ replicate + condition). The horizontal line indicates a significance level of padj < 0.05. Arrows indicate genes outside axis limits. Top-40 consistently changed genes between A and C are labelled (total overlapping: down = 22, up = 70). Fig 5D. Barplot of DESEQ normalized counts for *CEBPA* (Experiment as in B.) Fig 5E. Flow cytometry staining for CD15 and CD34 over time in cells overexpressing 4xPLDLS or 4xPLASS (constitutively expressed) from a p50MX retroviral vector. Timepoints indicate days since selection (on zeocin) is started.

## Discussion

This study demonstrates that *CEBPA*, a gene that is indispensable for neutrophil development, is directly repressed by MECOM in AML with 3q26 rearrangements through the +42kb *CEBPA* enhancer. Binding of Mecom to the +37kb murine homolog of the +42kb human enhancer has been reported in immortalized mouse bone marrow cells, but the effects on *Cebpa* transcription and a mechanism of action remained unclear^27^. Here we demonstrate that upon rapid MECOM depletion, *CEBPA* becomes upregulated, leading to substantial remodeling chromatin and changes in the transcriptome of inv(3) cells. Accordingly, overexpression of *CEBPA* alone can bypass the repression of differentiation by MECOM. A repressive effect of MECOM on *CEBPA* is further supported by the low *CEBPA* mRNA levels in 3q26-rearranged AMLs compared to other AML subtypes.

CEBPA is a master regulator of myeloid development and its dysregulation is a hallmark of several AML subtypes outside of inv(3)/t(3;3) (reviewed in: ^43,44^). Firstly, it can be directly mutated (usually an in-frame mutation in the bZIP domain in one allele and a frameshift mutation in the other allele), leading to overexpression of mutant *CEBPA*. Conversely, its locus can be silenced by DNA methylation in mixed lymphoid/myeloid phenotype leukemias ^40^. Low *CEBPA* expression can also result from repression at the +42kb enhancer. Fusion gene *RUNX1::RUNX1T1* (as a result of t(8;21)) and *RUNX1::MECOM (*t(3;21)) can repress *CEBPA* transcription via this enhancer ^42,45^. It has also been hypothesized that *CBFB::MYH11 (*as a consequence inv(16)) is able to repress *CEBPA* through an unknown RUNX1-dependent mechanism ^46^). In each of those three subtypes, transcription factor RUNX1 is either directly involved or required. In the case of 3q26-rearranged AML, binding of RUNX1 to the +42kb enhancer increases slightly upon MECOM depletion. Genome wide, sites that gain RUNX1 binding are enriched for CEBPA motifs. It would be interesting to investigate what the role of RUNX1 is in the regulation of *CEBPA* and what the interplay is between these three TFs in the context of 3q26-rearranged AML.

When deleting the +42kb enhancer, we were only able to derive heterozygous clones unable to differentiate upon MECOM depletion. Rather than showing a direct effect of MECOM binding at this enhancer, this demonstrates that the +42kb enhancer is required to upregulate *CEBPA* sufficiently in this context. The fact that this effect occurs in heterozygous clones, where CEBPA is only slightly upregulated upon MECOM knock-out, suggests that the effect of CEBPA is dose-dependent. Perhaps a positive autoregulatory feedback loop requires CEBPA to bind the 42kb enhancer to further upregulate its transcription.

We demonstrated that *CEBPA* overexpression bypasses the MECOM-mediated differentiation block (Figure 4D), but that this less effective than MECOM degradation (Figure 1C). This suggests that there are other MECOM targets that could play a role in this process. Indeed, a handful of other target genes apart from *CEBPA* were upregulated upon MECOM degradation. Although CEBPA was the only one of those which was consistently found as MECOM target across different experiments, it would be interesting to study whether overexpression of those genes either alone or in combination with *CEBPA* will bypass the MECOM-induced block of differentiation even more effectively in those leukemia cells.

The gain of *CEBPA* transcription upon MECOM degradation via the +42kb enhancers is accompanied by loss of CTBP2 binding. We recently demonstrated that interaction with CTBP1/2 is essential for leukemic transformation by MECOM, but it is unclear which MECOM cofactors are CTBP1/2-dependent^11^. Our results demonstrate that blocking of MECOM-CTBP2 interaction upregulates *CEBPA* levels and induces differentiation, but not to the same order of magnitude found when MECOM was degraded. This could be explained by the fact that, while 4xPLDLS treatment displaces CTBP from the chromatin while MECOM is still bound, potentially hindering the binding of activating transcription factors to the +42kb enhancer of *CEBPA*. We previously demonstrated that multiple repressor proteins interact with MECOM in AML cells. Therefore, it is also possible that multiple other interaction partners (CTBP-dependent and independent) play a role in gene repression by MECOM. CRISPR/Cas genetic screens, using gRNAs directed to genes encoding the factors present in the MECOM complex, should uncover whether any of those other co-factors are also involved in repressing *CEBPA* transcription.

The results presented here firmly establish that *CEBPA* is repressed by MECOM in MUTZ3 cells. In the past, MECOM has been reported to repress as well as activate target genes. In our SLAM-seq experiment, we find no evidence that MECOM directly activates target genes. In contrast, in experiments where MECOM is depleted for longer periods of time, we do identify downregulated genes, which most likely reflect secondary effects rather than direct activation by MECOM. In addition, it is conceivable that in different MECOM positive AML cell types MECOM regulates overlapping but also different target genes. This is supported by our observation that in *KMT2A*-rearranged MECOM+ patient samples, *CEBPA* levels were not decreased compared to the general AML population.

3q26/MECOM-rearranged leukemias have a severely adverse prognosis with current therapies. Given the critical role of MECOM in maintaining the leukemic phenotype, therapy resistance could be overcome by targeting this transcription factor. Our findings suggest that, when developing such targeted therapies, *CEBPA* levels can be used as a read-out for MECOM function in this context. Taking advantage of this property, we propose a two-step drug screening: first, in vitro assays could be employed to identify drugs that lift *CEBPA* repression, which could next be tested on their efficacy to block leukemic outgrowth of MECOM-rearranged AML cells in our previously developed xenotransplant models. Our studies demonstrate, for the first time, a strong mechanistic link between two key genes in hematopoietic development, *MECOM* and *CEBPA* and provides an explanation about how MECOM is able to maintain the stem cell state in AML with 3q26/MECOM rearrangements.

## Methods

### Cell lines

MUTZ3 (DSMZ, cat ACC295) cells were cultured in Alpha mem medium (Gibco cat 2571038) supplemented with 20% supernatant of urinary bladder carcinoma cell line 5637 (DSMZ, cat ACC35) ^48^, and 20% FCS (Sigma-Aldrich, cat F7524-500ml). Medium used was supplemented with 50U/ml penicillin and 50 U/ml streptomycin (Gibco, cat 15140122). All cell lines were routinely confirmed to be mycoplasma free by using the MycoAlert Mycoplasma Detection kit (Lonza, cat no LT07-318).

### Generation of viral supernatants

Lentiviruses were produced by co-transfecting 293T cells using psPAX2 (Addgene, cat 12260), pMD2.G (Addgene, 12259) and a lentiviral vector. Lentiviral supernatants were harvested at 72 HRS post transduction. All transfections were performed using Fugene-6 transfection reagent according manufacturers protocol (Promega, cat E2691).

### RT-PCR human and mouse MECOM

A minimal of 1×10^6^ cells were resuspended in 1ml Trizol (Life Technologies, cat 15596018) and RNA was isolated following manufactures protocol. 1 µg RNA was used in reverse transcription using Superscript II reverse transcriptase (Life Technologies, cat 18064-014). Human or mouse MECOM PCR was carried out using specific primer sets (see table) using a ABI7500 real-time PCR cycler. Expression was normalized using mouse HPRT or human PBGD.

### Western blot analysis

Cells were lysed in a buffer containing 20 mmol/L Tris-HCl, 138 mmol/L NaCl, 10 mmol/L EDTA, 50 mmol/L NaF, 1% Triton, 10% glycerol which was supplemented with protease inhibitors SigmaFast, Na_3_VO_4_ and reducing agent DTT (all chemicals purchased from Sigma-Aldrich). Protein content of lysates was measured by Pierce BCA protein Assay (Thermo scientific, cat 23227). 40 µg protein was loaded on a 4-12% Bis-Tris polyacrylamide gel (Thermofisher cat NP0321) and ran in a Mini Gel Tank (Thermofisher) in 1X MOPS buffer (Thermofisher cat NP001) Proteins were semi-dry blotted onto a 0.2 µM nitrocellulose membrane (Sigma cat GE10600001) and protein levels were detected by specific antibodies directed against: Human MECOM (Cell signaling, cat 2265), CTBP1 (BD, cat 612042), CTBP2 612044 (BD, 612044), V5 tag (Life Technologies, cat R96025), FLAG tag (Sigma-Aldrich, cat F3165), MYC tag (Santa Cruz, cat SC-40) CEBPA (CST cat. 8178), HA (Santa Cruz cat. SC805) and β-actin (Sigma-Aldrich, cat no A5441). Proteins were visualized using the Odyssey infrared imaging system (LI-COR Biosciences).

### Flow Cytometry

Flow cytometric analysis of mouse bone marrow cells or MUTZ3 cells was done with specific antibody staining using human CD34-PE-Cy7 (BD Biosciences, cat 348811), human CD15-APC (Sony, cat 2215035), mouse CD11B-APC (BD, cat 553311) Cells were analyzed on a FACSymphony (BD Bioscience). Flow data is analyzed with FlowJo. Stacked area plots are made in ggplot. Flow data is visualized in FlowJo, or CytoML and ggCyto for stacked histograms.

### shRNA knockdown

EVI1 was downregulated by transducing MUTZ3 cells using Mission shRNA’s directed against human EVI1 (Sigma Aldrich). At 48HRS post transduction cells were selected on 1 µg/ml Puromycin. Knockdown of EVI1 was confirmed by Western blot and RNA was harvested at 24,48,72,96 120HRS of Puromycin selection. 700 ng total RNA was used in library preparation using the KAPA RNA HyperPrep kit with RiboErase (Roche, cat KK8561). Libraries were paired-end sequenced (2 × 100 bp) on the Novaseq 6000 platform

### Models: MUTZ3 MECOM-V5-AID-T2A-GFP OsTIR1

Fragment containing a spacer with AID-V5-tag was ordered as a gblock (IDT) and cloned into pJet (Thermofischer, cat K1231) by blunt-end cloning. The G-block and repair template generated for a MUTZ3-MECOM-T2A-GFP cell line previously published ^10^ were PCR amplified and cloned as new repair template using Gibson assembly (NEB, cat E5510S). The PAM sequence (sgRNA AGCCACGTATGACGTTATCA) was omitted in the repair template. Guide RNA was produced using HiScribe™ T7 High Yield RNA Synthesis Kit (NEB, cat E2040S; 320ng guide, 20ug Cas9, 10 ug donor template in 100 uL transfection volume). Cells were nucleofected with sgRNA, Cas9 (IDT, cat 1081061) and the repair template using the Neon (Thermofisher) with buffer R and settings at 1500V, 20ms, 1 pulse for MUZT3 or 1350V. GFP positive cells were sorted using a FACS Aria III cell sorter. After 3 serial sorts GFP positive cells were single-cell sorted into a round-bottom 96-well plate. Clones were tested for integration with V5 protein expression with western blot. Correct clones were used in transduction with OsTIR1 3xmyc T2A puro. Cells were placed under puromycin selection and were single-cell sorted. Clones were screened for expression of the MYC-tag on western blot. MECOM degradation was confirmed in OsTIR1 expressing clones by western blot for MECOM or V5 tag after exposing cells to 500 µM Auxin for indicated timepoints (Sigma-Aldrich, 15148-2g).

### Cloning lentiviral CRISPR guides

To delete 700bp of the +42kb CEBPA enhancer, an upstream (U6 – CTCTATCCCTCTACCCAGAG) and downstream (H1 – GAGCAAATCACGAAGCCCAG) sgRNA was cloned in a lentiviral dual guide expressing vector pDecko-mCherry-Puro (Addgene #78534)^49^. Briefly, the pDecko vector was digested with BsmBI and 6 assembly guides containing the guides separated by an internal BsmBI site which was assembled with Gibson assembly (NEB, as above). The molar ratio of each oligo to backbone was 32:1 (with 180 ng backbone). This intermediate vector is digested again with BsmBI and the H1 internal promoter is PCR amplified from the original vector and assembled with Gibson assembly to create the final vector (3:1 insert:vector). For knock-out of *MECOM* (MECOM.4) and mutations in the *GATA2* enhancer driving *MECOM* expression (ENH3, ENH11, ENH8), a single guide was cloned into lentiviral expression backbone pLentiV2 U6-IT-mPgk-mCherry using BbsI digestion as described before^10^. Guide sequences are included in Supplementary Table 1. Guides were lentivirally transduced (see above for lentivirus production) into a previously generated MUTZ3-iCas9 clone with doxycycline-inducible expression of Cas9^10^. Cas9 expression was induced with 1 ug/mL doxycycline (Takara Cat 631311). For the deletion of the *CEBPA* enhancer, mCherry+ cells were sorted and doxycline-induced for 7 days. 4 days later, clones were derived using limiting dilution which were tested for the deletion of the +42kb enhancer by PCR (1212 bp = WT; 381 bp = deletion; See Supplementary table 1 for sequences).

### *CEBPA* overexpression

To overexpress CEBPA, an empty pCW57-BLAST lentiviral backbone with a doxycycline-inducible promoter (Addgene #80921) was digested with NheI and BamHI. A Gblock was ordered (IDT) containing NheI-CEBPA cDNA – 3x HA – MluI-stop-BamHI in a pUC57 vector (pUC57-gblock-CEBPA-3xHA). The Gblock was also digested with NheI and BamHI. The backbone was dephosphorylated with AP (NEB #M0289S) and the insert phosophorylated with PNK (NEB #M0201S) and fragments assembled with a DNA ligation reaction at a 3:1 insert:backbone ratio with T4 DNA ligase (NEB #M0202S). MUTZ3 cells were transduced with lentivirus (see above) and selected on blastcidin (12 ug/mL; ThermoFisher cat 210-01) for 9 days.

### SLAM-seq

MUTZ3 MECOM V5 AID GFP OsTIR1 expressing cells were treated with 500 µM/ml Auxin (Sigma-Aldrich, 15148-2g). After 1.5, hours nascent RNA was labeled for 3 hours using 300 µM/ml 4-Thiouridine (Lexogen, cat 061.24). At 3 HRS of 4-Thiouridine labeling RNA was isolated using Qiagen rnease mini kit (Qiagen, cat 74104). Alkylation of thiol groups was performed according to manufacturer’s protocol (Lexogen, cat 061.24). For RNA sequencing, 500 ng total RNA was used to generate libraries using the KAPA RNA HyperPrep kit with RiboErase (Roche, cat KK8561). Amplified sample libraries were paired-end sequenced (2 × 100 bp) on the Novaseq 6000 platform.

### ChIP-seq

25×10^6^ cells derived from cell lines or patient material were used in ChIP. Cells were pelleted and resuspended in PBS containing 2 mM disuccinimidyl glutarate. Cells were incubated for 45 minutes at RT. Cells were pelleted and resuspended in PBS containing 1% formaldehyde and incubated 15 minutes RT. Crosslinked cells were pelleted and washed 3x using lysis buffer (10 mM Tris HCL pH 7.5, 10 mM NaCl, 3 mM MgCl2 and 0.5% NP40). In between washing steps samples were incubated on ice for 5 minutes. Next, Nuclei were lysed using nuclei lysing buffer containing 160 mM NaCl, 0.8% SDS, 5 mM Sodium Butyrate, 1/5 pre IP buffer (10 mM Tris HCL pH 7.5, 10 mM NaCl, 3 mM MgCl2, 1 mM MgCl2, 4% NP40), 1× Roche protease inhibitor and 10 mM phenylmethylsulfonyl fluoride. Lysates were sonicated on a Biorupter (Diagenode) and size was evaluated. Chromatin was diluted at least 5× using IP dilution buffer (Upstate), 5 µg antibody and 30 µl protein G beads were added. Samples were incubated overnight while mixing. Next day, beads were washed according the upstate protocol. DNA was eluted, decrosslinked and quantified. 10 ng ChIP DNA was used in library preparation using MicroPlex library preparation kit (Diagenode, cat C05010013). Amplified sample libraries were paired-end sequenced (2 × 100 bp) on the Novaseq 6000. The following antibodies were used in ChIP-seq: H3K27AC (Diagenode cat C15410196, lot A1723-0041D), RUNX1 (Abcam cat AB23980, lot GR3364239-1), P300 (Diagenode cat C15200211, lot 001), H3K9ME3 (Diagenode, cat no C15410193, lot A2217P), H3K27ME3 (Diagenode cat no C15410195, lot A0824DD), CTBP2 (BD cat no 612044, lot7069947) and V5 (ThermoFisher #R960-25). Track for EVI1 in primary material was published previously ^11^

### 4C-seq (Circular Chromosome Conformation Capture Sequencing)

10 x10^6^ cells were collected and chromatin was cross-linked using 2% formaldehyde in PBS with 10% FCS for 10 min at room temperature. Glycine (0.125M) was added to quench the crosslinking reaction. Cells were centrifuged and nuclei were isolated by resuspending the cells in fresh prepared nuclei lysis buffer (50 mM Tris-HCl pH 7.5, 0.5% NP-40, 1% Triton X-100, 150 mM NaCl, 5 mM EDTA and 1× protease inhibitors). Nuclei were permeabilized in 1.2x primary restriction enzyme buffer with 0.3% SDS for 1 hour at 37°C, followed by 1 hour at 37°C with 2.5% Triton X-100. Cross-linked DNA was digested overnight with DpnII (the primary restriction enzyme; 10,000 units/ml, New England Biolabs, R0543), followed by proximity ligation overnight at 16°C. Digested DNA was diluted in 7 mL 1× ligation buffer (66 mM Tris-HCl, pH 7.5, 5LmM MgCl_2_, 5 mM DTT, 1 mM ATP) and 50 U T4 DNA Ligase (Roche cat 10799009001) were added. Cross-links were removed at 65 °C using 30 ug prot K. DNA was purified with phenol extraction followed by ethanol precipitation. The purified DNA was digested with NlaIII (secondary restriction enzyme; 10,000 units/ml, New England Biolabs cat R0125), followed again by proximity ligation overnight at 16°C. Digested DNA was diluted to 5ug/ml in 1× ligation buffer and 50 U T4 DNA Ligase was added. The circular ligated 4C DNA template was precipitated, concentrated and purified using Qiaquick PCR purification kit (Qiagen cat 28104). The library preparation of the 4C DNA template involves two PCR steps. In the first PCR step, 4 reactions with 200 ng 4C template were performed to amplify the fragments ligated to the viewpoint using Expand Long Template Polymerase kit (Roche cat 11681834001; reaction conditions: 5uM reading primer, 5uM non reading primer; PCR programme: 94 °C for 2 min, 16 cycles 94 °C for 10 s; 55 °C for 1 min; 68 °C for 3 min and 68 °C for 5 min). All PCR reactions were pooled and purified with 0.8x AMPure XP beads (Beckman Coulter A63881). In the second PCR step with the same kit, 0.25 uM universal primers containing the Illumina adapters, a 6-nt index sequence and the Illumina sequencing primer sequences was used (PCR programme: 94 °C for 2 min, 16 cycles 94 °C for 10 s; 60 °C for 1 min; 68 °C for 3 min and 68 °C for 5 min). The amplified 4C DNA library prep was purified using Qiaquick PCR purification kit and subjected to next-generation sequencing on the IIlumina NovaSeq 6000 platform.

### ATAC-seq

MUTZ3 MECOM V5 AID GFP OsTIR1 expressing cells were treated with 500 µM/ml Auxin. At 4 and 24HRS cells were prepared for ATAC sequencing. Cells were pelleted and washed using PBS. Next, nuclei were isolated by lysing cells in 1mL lysis buffer (0.3M sucrose, 10 mM Tris HCl pH 7.5, 60 mM KCl, 15 mM NaCl, 5 mM MgCl, 0.1 mM EGTA, 0.1% NP40, 0.15 mM Spermine, 0.5M Spermidine and 2mM 6-A). Lysates were incubated on ice 5 minutes. Nuclei were pelleted and resuspended in 50 µl transpose mix containing 2.5 µl TD1 transposase (Illumina Cat FC-121-1030). Samples were incubated 30 minutes at 37°C. DNA was purified using Qiagen’s PCR min elute kit (Qiagen cat 28004). DNA was amplified using indexed primers. Amplified sample libraries were paired-end sequenced (2 × 100 bp) on the Novaseq 6000.

### RNA-seq Patients

For RNA-seq of patient samples, previously generated datasets in our department were combined by selecting samples based on the same sequencing method (libraries prepared by KAPA RNA HyperPrep and paired-end sequenced (2 × 100 bp) on the Novaseq 6000 as above). Transcript quantification was performed with Salmon^50^ and the settings --validateMappings --gcBias --seqBias, using the Homo sapiens hg38 reference genome and the refGene transcriptome database ^6,38–40^. Patients provided written informed consent in accordance with the declaration of Helsinki at the centre where the material was collected.

## Data availability

Raw data will be deposited on Zenodo with DOI 10.5281/zenodo.14505323. Patient sequencing data will be deposited at EGA under identifier EGAS00001008005 and raw data is available under a data transfer agreement.

### SLAM-seq

Analysis of SLAM-seq data was based on <u>SLAM-DUNK</u> ^51^. The slamdunk pipeline was adapted to take total RNA as input instead of 3’UTR only. Differential gene expression analysis was performed using the R package DEseq2 and used counts from nascent RNA only as based on the SLAM-DUNK pipeline. Briefly, genes with no T>C-converted counts were excluded. DESeq2 was run on the T>C-converted counts with size factors estimated from the total counts. As a design formula, ∼ clone + condition was used to correct for baseline differences between clones. ApeGLM was used for fold-change shrinkage of the auxin versus no_auxin contrast.

### Total RNAseq

Transcript abundance was quantified from RNA-seq fastq files directly with Salmon (v0.13.1) in pseudoalignment mode and the settings --validateMappings --gcBias --seqBias, using the Homo sapiens hg38 reference genome and the refGene transcriptome database Transcript-level counts were aggregated to genes with tximport (v1.26.1), and differential gene expression analysis was performed with DEseq2 (v1.44.0). A significance cutoff of log2 FC > 1 and adjusted P-value< 0.05 was used to determine the set of DE genes.

### shRNA knockdown analysis

*CEBPA* genesets were downloaded from <u>mSigDB</u> and <u>Enrichr</u>. Fgsea was used for gene set enrichment. Genes in the leading edge at any timepoint for the CEBPA UP target set from Tavor et al^47^ (included in mSigDB) were extracted and filtered for significant DE expression at any of the 5 time points. Genes were sorted by the first timepoint where they were detected as significantly differentially expressed. To generate the heatmap, the mean of the control samples at the given timepoint were subtracted from all values; the resulting values were row-normalised with a z-score. The heatmaps were generated separately for each shRNA.

### Patients RNAseq analysis

Patients with known silencing of the *CEBPA* locus were excluded. *CEBPA* bZIP mutants were grouped into a separate category as it is known their *CEBPA* expression is higher and this could increase the mean of the other_AML group. Patients were excluded from the analysis if they had known silencing (n = 2) or a deletion (n = 1) of the entire *CEBPA* locus. KMT2A status was based on karyotyping, PCR and fusion gene detection in RNA-seq. 3q26 status was determined by karyotyping or 3q-capture sequencing. Inv(16) and t(8;21) was determined based on karyotyping.

### ChIP/ATAC-seq

ChIP-seq reads were aligned to the human reference genome build hg19 with bowtie (v1.1.1) ^52^. For statistical analysis, the R package DiffBind (v 3.14.0) was used. For DiffBind, peaks were called with MACS2 (v2.2.7.1) with default parameters and a matched input file (same ChIP protocol on the same cell type) for transcription factors. For ATAC-seq, the same default settings were used but no input file. The –broad argument was used for peak calling on histone marks. For H3K27ac, peaks were centered on ATAC signal in any sample prior to analysis. For this, first, H3K27Ac consensus peaks were determined based on broad peak calls with DiffBind and summits = FALSE. This consensus set was overlayed with the summits from the ATAC consensus peakset (derived by DESeq2 with summits = TRUE). The ATAC peaks that were inside a H3K27Ac consensus peak were selected and widened to 2000 bp for DiffBind analysis of H3K27ac signal. This ensured symmetrical H3K27ac windows centered on the internal ATAC peak and was done to improve motif enrichment analysis. Intra-sample group variance was inspected with PCA and cluster heatmaps and default DiffBind settings were used for normalization. To annotate significant peaks, for each factor, peaks were annotated to promoters using GenomicFeatures in R. To link CREs to putative target genes, peaks were lifted to hg38 and the public dataset with ChIPseq marks and Hi-C for progenitor subsets was used ^53^. The top-15 CRE peaks for each factor were manually annotated in this way (peaks that were not included in a loop were not shown; this was in particular for the RUNX1 ChIP-seq). A consensus annotation set consisting of any promoter significant at any timepoint, and the top-15 manually annotated peaks were used to annotate all ChIPseq/ATAC-seq volcano plots.

For ChIP-seq heatmaps (tornado plots), the R package Genomation was used. Peaks (consensus peaks determined by DiffBind) were sorted on either MECOM or CTBP2 enrichment and widened symmetrically to 2000 bp. These reference peaks were then used as windows to count signal in the BAM files of all tracks shown. The heatmaps are winsorised (“overexposed”) on (0,99), meaning all signal above the 99^th^ percentile of peaks are the same color. For visualisation of specific genomic regions, the R package Gviz was used. Transcription factor motif enrichment was performed with AME from the MEME suite using the memer package in r (v1.12.0). Peaks for each factor were divided in up (fold > 0 & FDR < 0.05), down (fold < 0 & FDR < 0.05) and unchanged (FDR ≥ 0.05). runAME was used on 400 bp peaks. For ATAC-seq footprinting analysis, reads for each replicate were combined into a single file (4hrs +auxin, 24hrs +auxin and control) and TOBIAS was used with default settings ^54^ (comparisons: 4 HRS vs control, 24HRS vs Control and 24HRS vs 4HRS).

## Acknowledgements

We would like to thank Remco Hoogenboezem for processing of sequencing data, Emma Boertjes and Peter Valk for help with annotating CEBPA mutations in patient data and Michael Vermeulen for FACS sorting.

## Author contributions

Conceptualization: R.D., E.B.

Data curation: R.M.L

Formal analysis: D.P., R.M.L.

Funding acquisition: R.D.

Investigation and validation: M.H., D.P., L.S., C.E.V., S.v.H., M.A., T.G., S.O., E.B.

Methodology: M.H., L.S., J.Z, E.B.

Resources: T.H., C.H, J.Z., B.W., B.B.

Software: D.P., R.M.L.

Supervision & Project administration: R.D.

Visualization: D.P.

Writing – original draft: D.P., M.H.

Writing – review & editing: D.P., R.D.

## Supplemental figures

**Figure S1:**
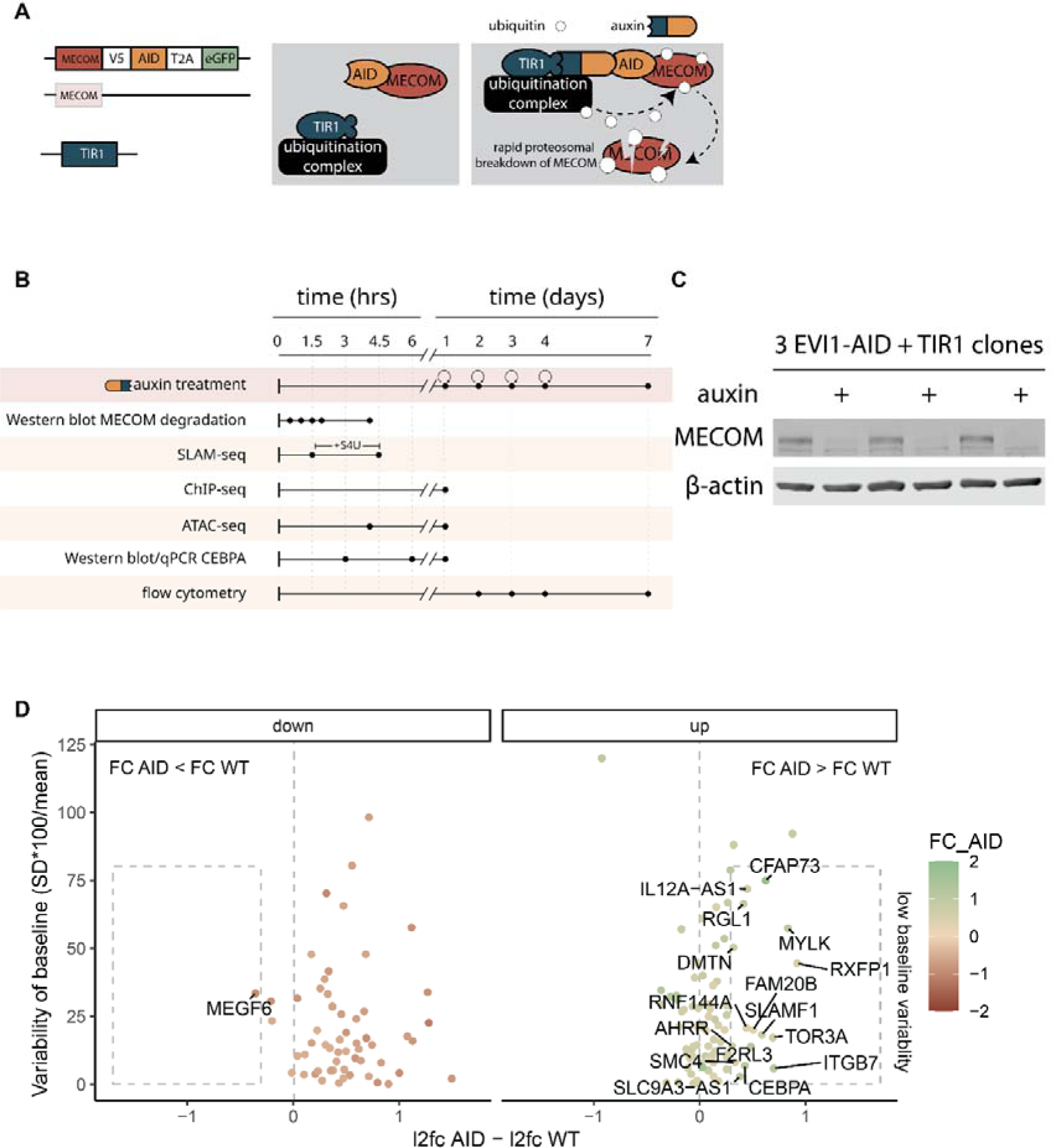
Supplement to Figure 1 Part 1. Fig S1A. Schematic depiction of the AID tag, followed by V5T2A-eGFP, introduced 3’ of MECOM in MUTZ3 cells (left). OsTIR1 was transduced into a MECOM-AID-V5-T2A-eGFP expressing clone (middle). Treatment of auxin leads to co-localization of MECOM-AID to the nuclear ubiquitination complex via OsTIR1 leading to rapid degradation of MECOM (right). Fig S1B. Timeline of all experiments on MUTZ3-MECOM-V5-AID auxin experiments. For multi-day experiments, auxin is refreshed daily as indicated by circular arrows. Fig S1C. Western blot analysis of 3 MUTZ3 MECOM-AID-TIR1 clones for MECOM (upper panel) and B-actin (lower panel). Cells were treated with 500 µM of auxin for 1.5 hrs. Fig S1D. Scatterplot of log2 fold-change (FC) in AID clones versus WT MUTZ3 (auxin/no auxin) versus the variability of baseline no_auxin measurements. Genes with low baseline variability and a more extreme fold-change in AID clones versus WT are highlighted in the rectangle.

**Figure S2:**
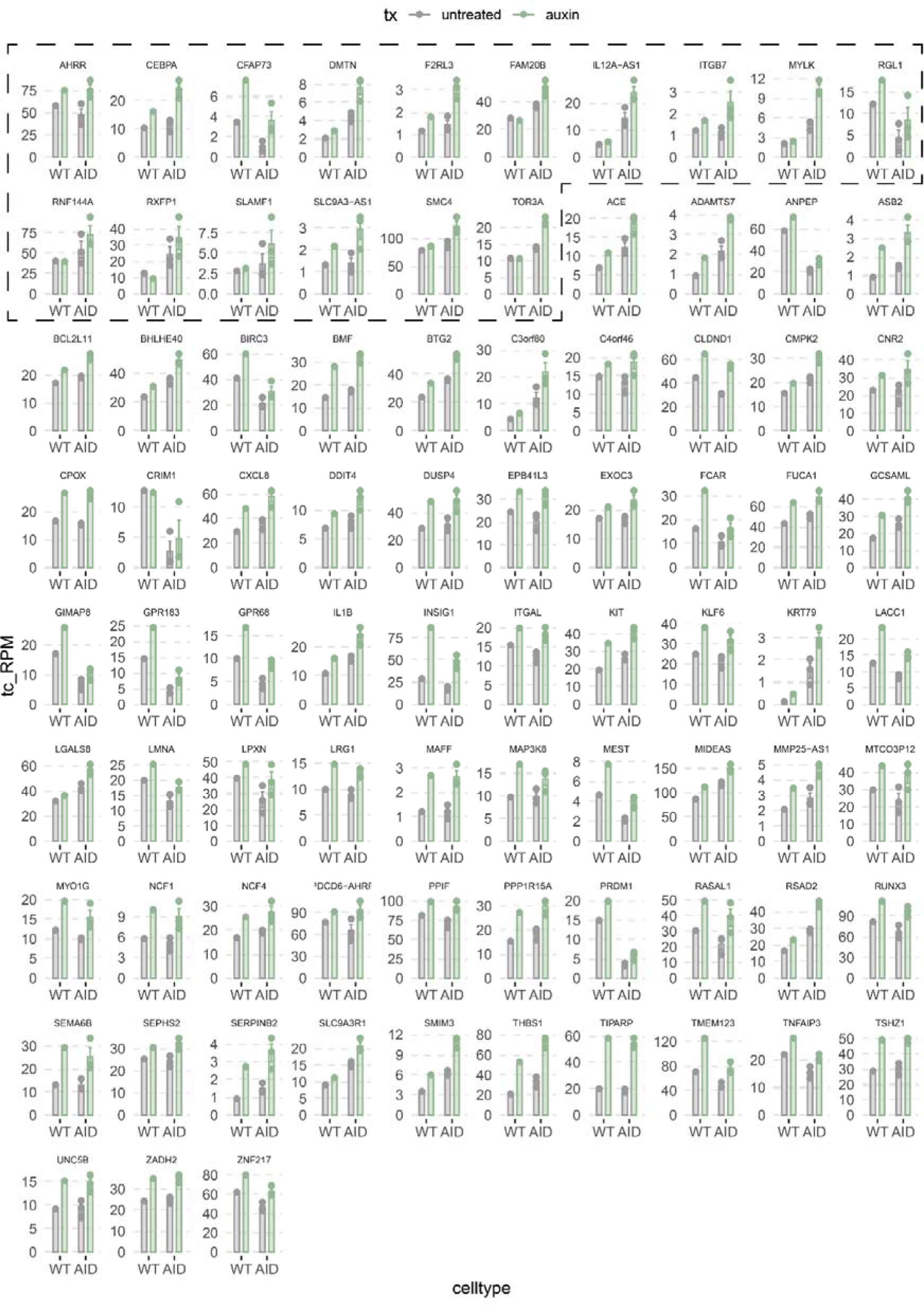
Supplement to figure 1 Part 2. Fig. S2A. Barplots of T>C converted counts of all upregulated genes (AID clones: auxin vs untreated) with the WT MUTZ3 cells included. Genes that were filtered for a higher fold-change in AID clones than WT are highlighted in the dashed box.

**Figure S3:**
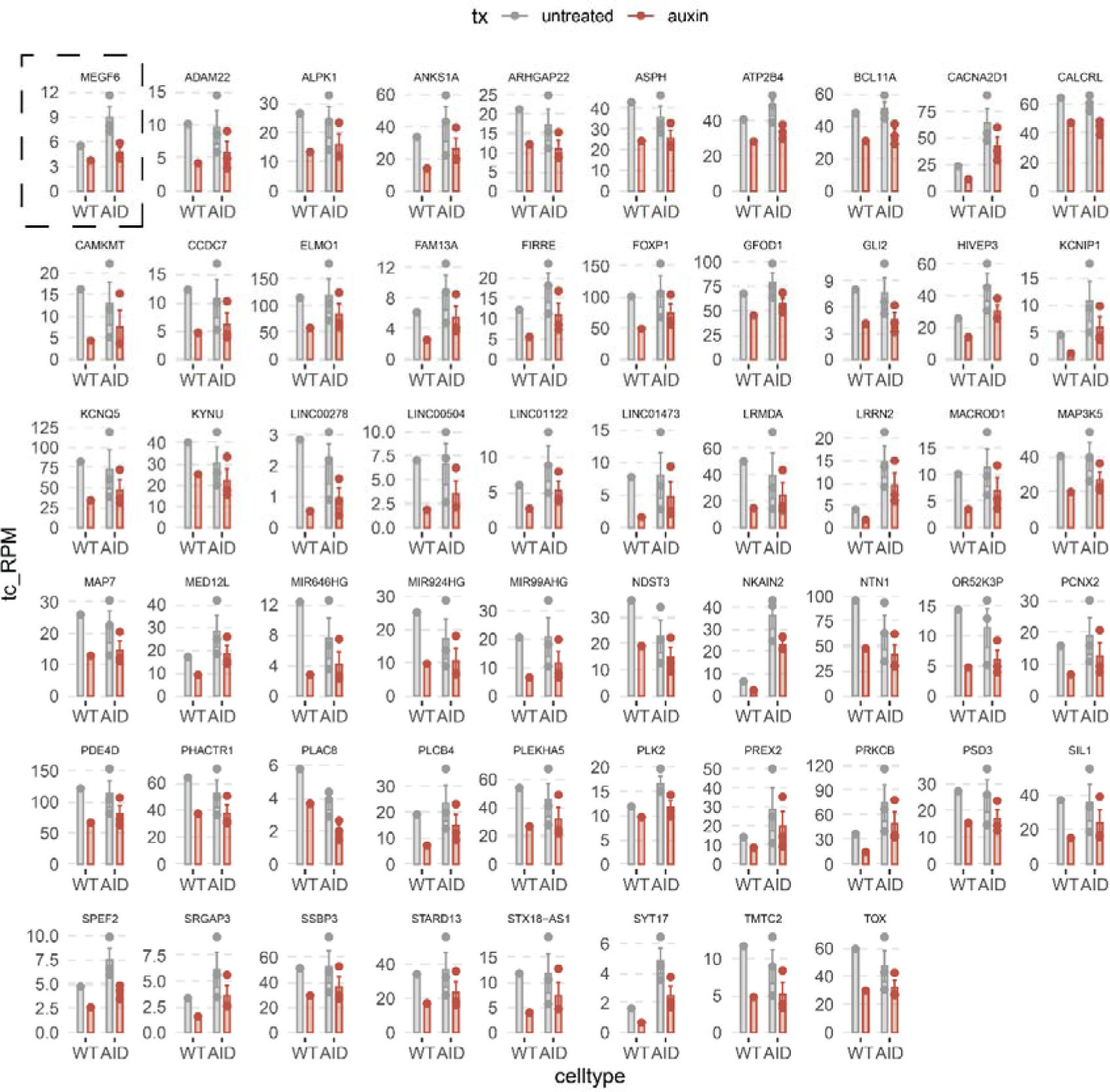
Supplement to Fig.1 Part 3. Fig S3A. Barplots of T>C converted counts of all downregulated genes (AID clones: auxin vs untreated) with the WT MUTZ3 cells included. Genes that were filtered for a higher fold-change in AID clones than WT are highlighted in the dashed box.

**Figure S4:**
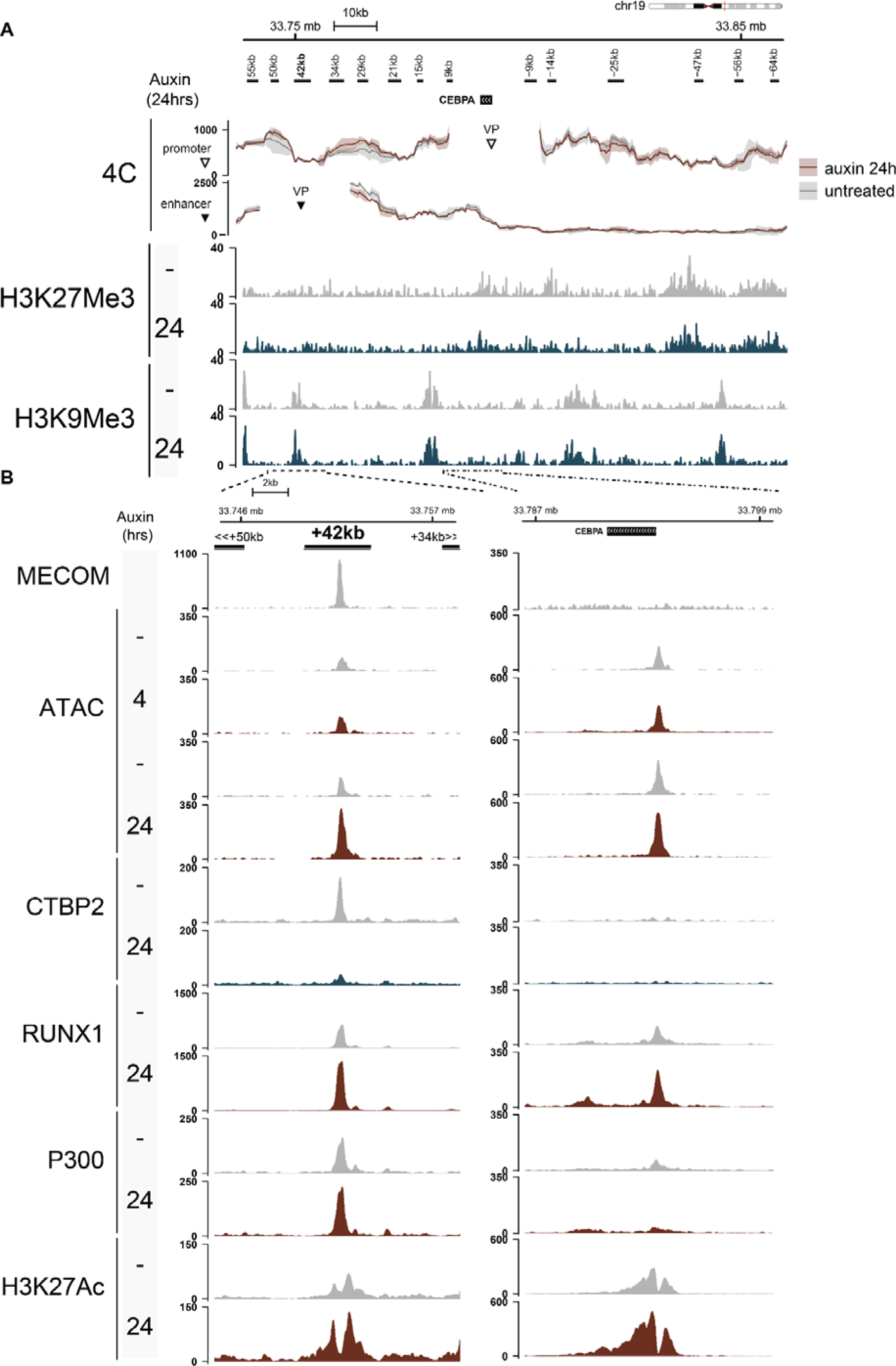
Supplement to Figure 2. Fig S4A. ChIP-seq tracks for indicated marks and 4C contact profiles in the *CEBPA* locus. **4C:** RPM-normalized data, viewpoint from the *CEBPA* promoter (top panel) or the +42kb enhancer (bottom panel). Mean + SD of two replicates from two clones is plotted. **ChIP-seq:** RPKM normalized data is plotted in hg19 coordinates. Fig S4B. Zoom-in on the *CEBPA* +42kb enhancer and the *CEBPA* gene body with all tracks shown in Fig. 2C.

**Figure S5:**
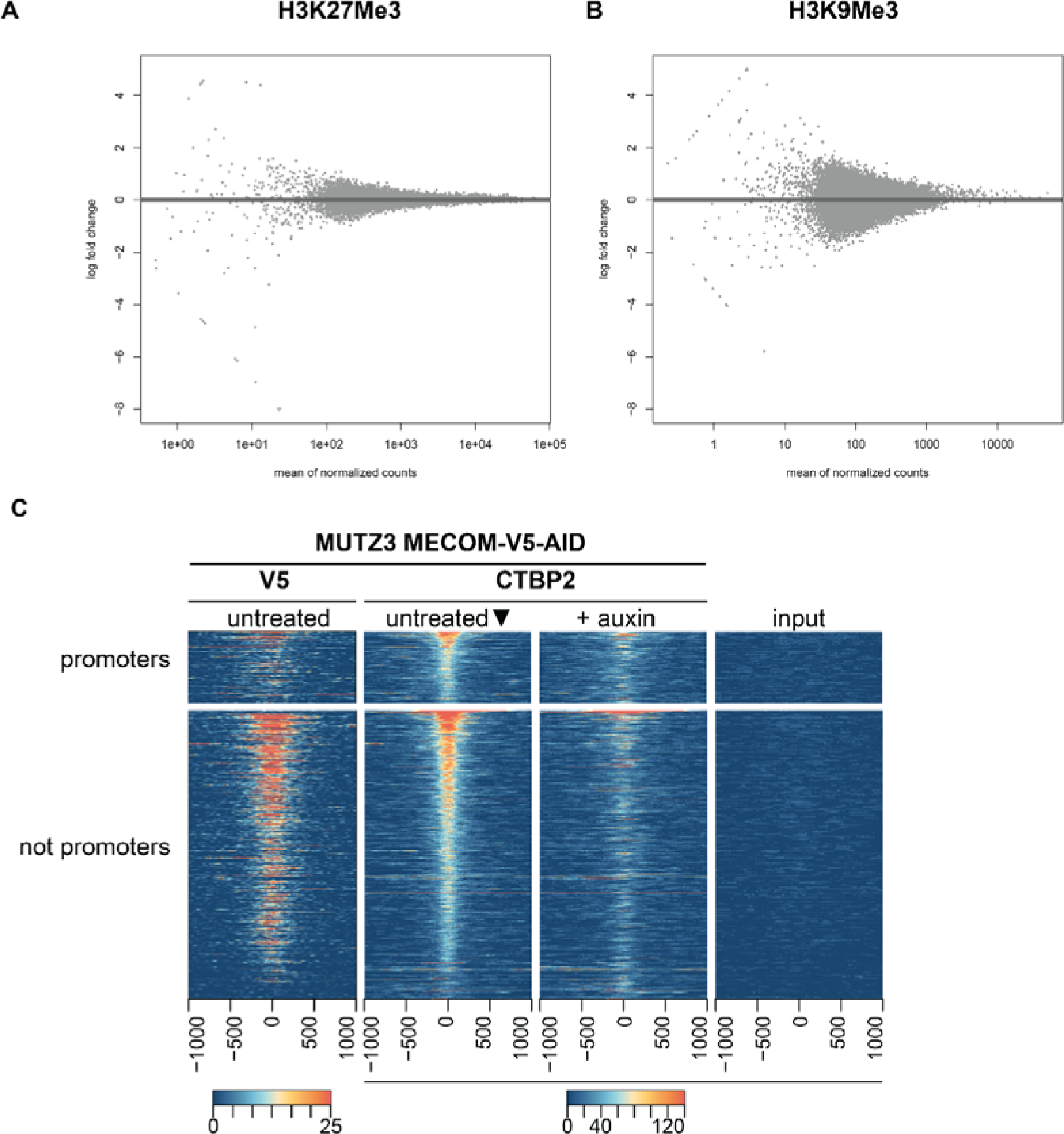
Supplement to Fig.2 Part 2. Fig. S5A. MA-plot of DESeq2 DiffBind analysis of counts for H3K9Me3. Fold-changes (Auxin vs. no auxin) are not shrunken. Fig. S5B. MA-plot of DESeq2 DiffBind analysis of counts for H3K27ME3. Fold-changes (Auxin vs. no auxin) are not shrunken. Fig. S5C. ChIP-seq heatmap of MECOM (V5) and CTBP2 in MECOM-AID-V5 clones, ranked by normalised CTBP2 binding as calculated by DiffBind in untreated control and subsetted according to whether they overlap promoters (-500/+2000bp of TSS) prior to widening to 2000 bp.

**Figure S6:**
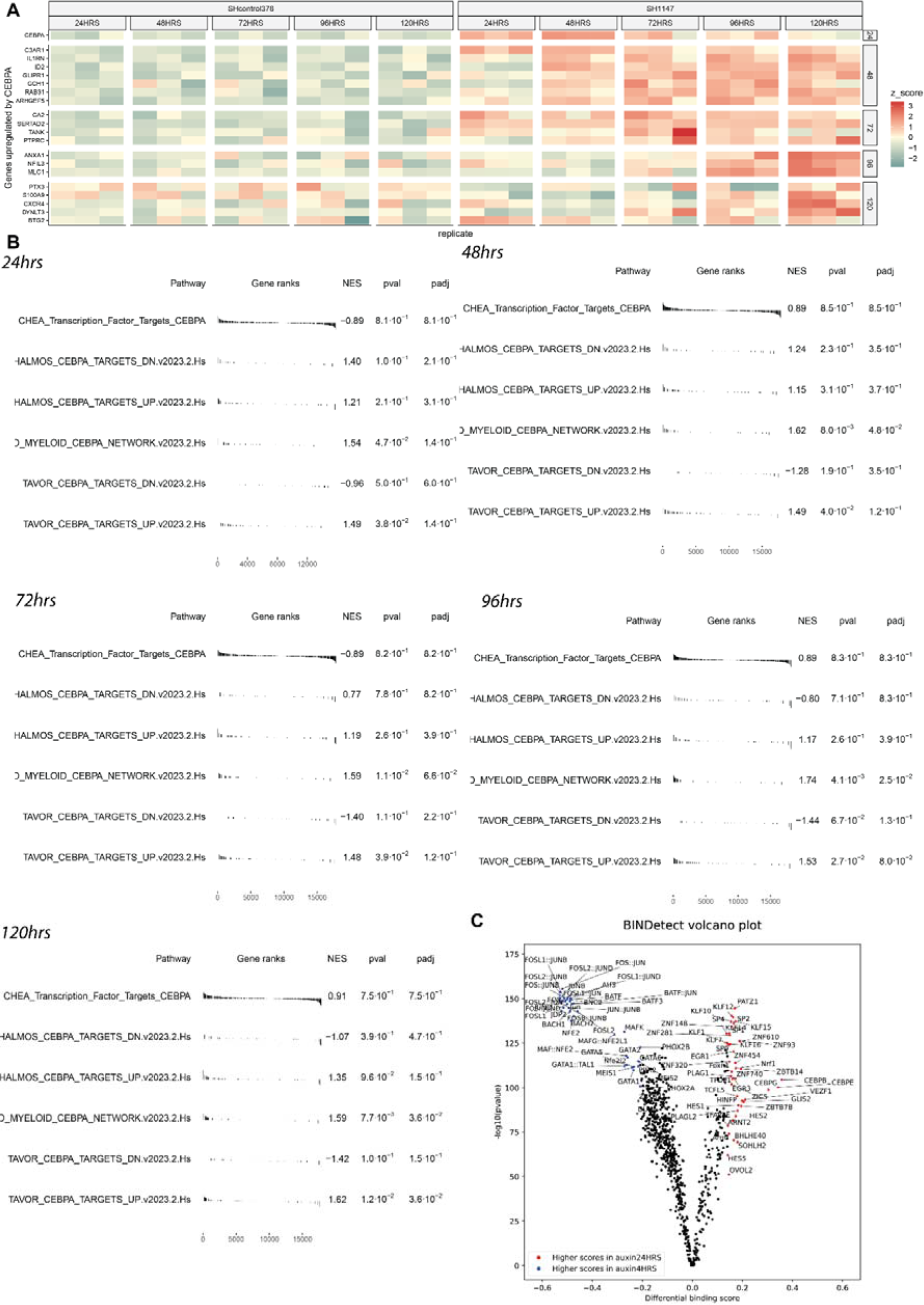
Supplement to Fig 3 Pt.1. Fig. S6A. Heatmap of z-scores of RNA-seq rlog-transformed counts of MUTZ3 cells treated with control shRNA or MECOM shRNA. Genes are selected from a geneset indicating genes upregulated by CEBPA (Tavor et al.). Genes are grouped according to when they are first upregulated post MECOM k.d. From rlog transformed counts, the mean of the control samples is subtracted prior to z-score calculation. Fig. S6B. Gene set enrichment computed by FGSEA for CEBPA related-genesets as described in the methods. Genes are ranked by -log10(padj) × sign(Fold-change). Fig. S6C. Volcano plot of ATAC-seq footprinting of MUTZ3-V5-AID-OsTIR1 cells after 24 hrs of auxin depletion (n = 2) compared to 4 hrs of depletion (n = 2) calculated with TOBIAS. Each dot is a single motif.

**Figure S7.**
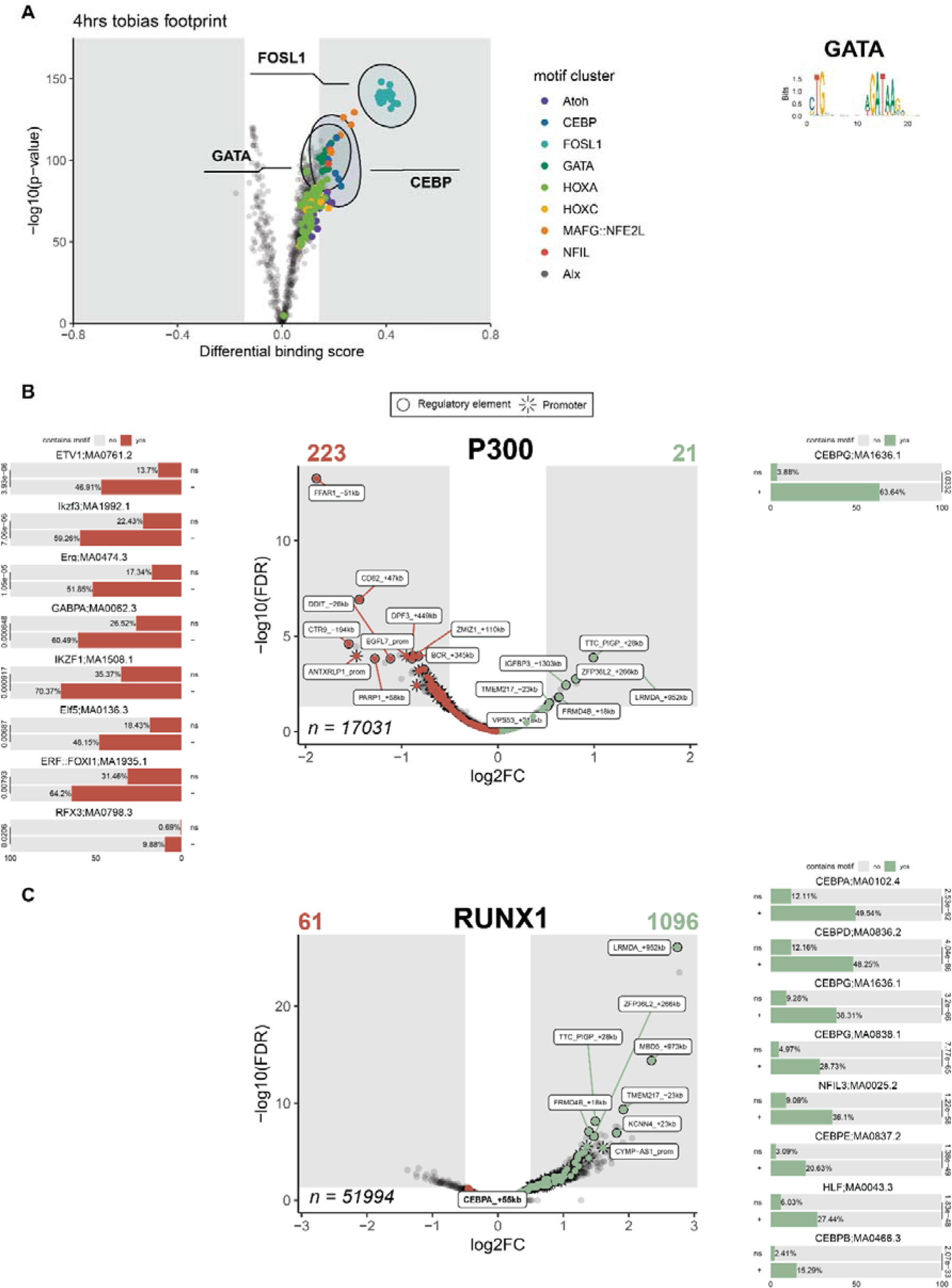
Supplement to Fig. 3. Fig S7A. Volcano plot of ATAC-seq footprinting of on MUTZ3-V5-AID-OsTIR1 cells after 4 hrs of auxin depletion (n = 2) compared to untreated control (n = 4), as calculated with TOBIAS. Each dot is a single motif, and motifs are colored by their cluster. Composite motifs of the indicated motif clusters are shown in the plot. Composite GATA motif for panel is shown on the right-hand side. Fig S7B. Volcano plot differential P300 binding (ChIP-seq) in MUTZ3-V5-AID-OsTIR1 cells after 24 hrs of auxin depletion versus untreated control (n = 2). Details of peak annotation in the Methods. AME motif enrichment is shown on the corresponding side of the plot. Bars represent the fraction of peaks that contain a high-score motif per set; significance is indicated with AME e-value across the bars. Top-8 motifs are shown per ChIP at an e-value cut-off is 0.05. All labels containing “CEBPA” at FDR < 0.05 are labelled in bold. Fig S7C. Volcano plot of differential RUNX1 binding (ChIP-seq) in MUTZ3-V5-AID-OsTIR1 cells after 24 hrs of auxin depletion versus untreated control (n = 2). Peak annotation is described in the methods. AME motif enrichment is shown on the corresponding side of the plot. Bars represent the fraction of peaks that contain a high-score motif per set; significance is indicated with AME e-value across the bars. Top-8 motifs are shown per ChIP at an e-value cut-off is 0.05. All labels containing “CEBPA” at FDR < 0.05 are labelled in bold.

**Figure S8:**
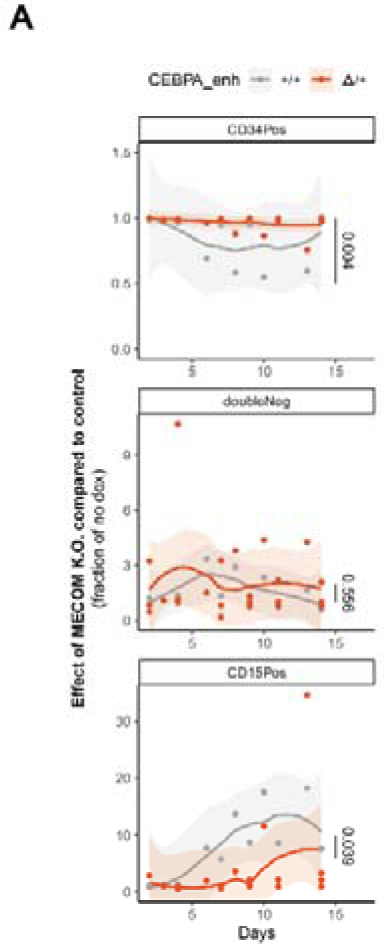
Sup to Fig 4. Fig. S8A. Summary of experiment in panel 3B on 4 heterozygous *CEBPA* enhancer deleted clones versus 2 bulk WT clones. Significance of the differences between WT and heterozygous deleted clones is calculated with anova_ez from the AFEX package with days as covariate. P-value shown on graph.

**Figure S9:**
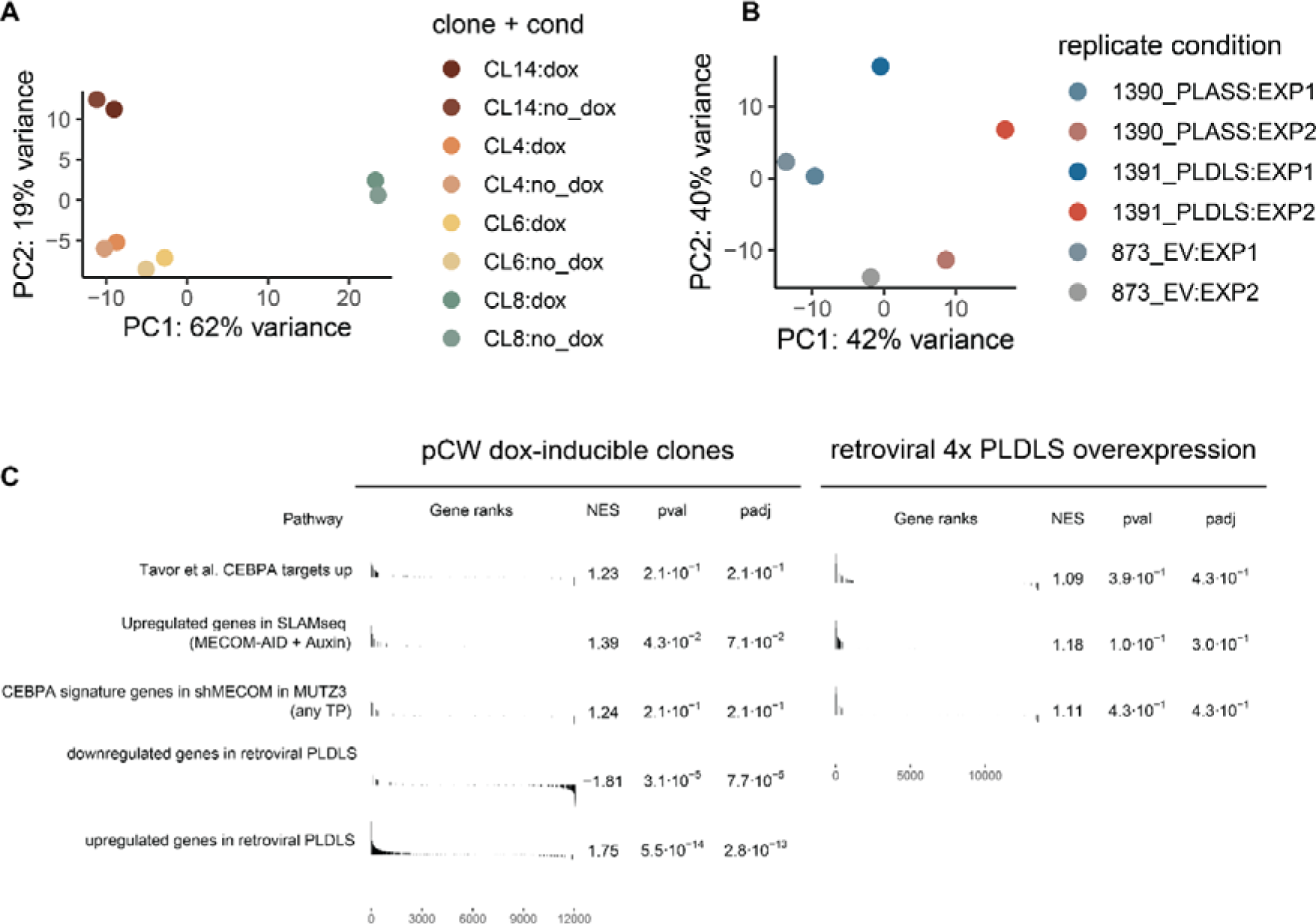
Supplement to Fig 5. Fig S9A. PCA of RNA-seq data of MUTZ3 cells transduced with pCW-4xPLDLS (dox-inducible). 4 individual clones are derived and dox-treated for 72hrs Fig S9B. PCA of RNA-seq data of MUTZ3 cells transduced with retroviral p50MX-4XPLDLS 72hrs after zeocin selection. Fig S9C. GSEA of MUTZ3 either transduced with pCW-4xPLDLS(left) or with retroviral p50MX-4XPLDLS (right). Genes identified as targets of CEBPA^47^, the CEBPA signature in the shRNA KD experiment and upregulated genes measured by SLAM-seq after MECOM degradation are included as gene sets. In addition, DE genes from the retroviral 4xPLDLS experiment are included at the bottom for the lentiviral system (pCW-4xPLDLS) only.

